# Investigating the Unbinding of Muscarinic Antagonists from the Muscarinic 3 Receptor

**DOI:** 10.1101/2023.01.03.522558

**Authors:** Pedro J. Buigues, Sascha Gehrke, Magd Badaoui, Gaurav Mandana, Tianyun Qi, Giovanni Bottegoni, Edina Rosta

**Author notes:** If an author’s address is different than the one given in the affiliation line, this information may be included here.

## Abstract

Patient symptom relief is often heavily influenced by the residence time of the inhibitor-target complex. For the human muscarinic receptor 3 (hMR3), tiotropium is a long-acting bronchodylator used in conditions such as asthma or chronic obstructive pulmonary disease (COPD). The mechanistic insights of this inhibitor remain unclear, specifically, elucidation of the main factors determining the unbinding rates could help develop the next generation of antimuscarinic agents. Using our novel unbinding algorithm, we were able to investigate ligand dissociation from hMR3. The unbinding paths of tiotropium and two of its analogues, N-methylscopolamin and homatropine methylbromide show a consistent qualitative mechanism and allowed us to identify the structural bottleneck of the process. Furthermore, our machine learning-based analysis identified key roles of the ECL2/TM5 junction involved at the transition state. Additionally, our results point at relevant changes at the intracellular end of the TM6 helix leading to the ICL3 kinase domain, highlighting the closest residue L482. This residue is located right between two main protein binding sites involved in signal transduction for hMR3’s activation and regulation. We also highlight key pharmacophores of tiotropium that play determining roles in the unbinding kinetics and could aid towards drug design and lead optimization.

**Description:** 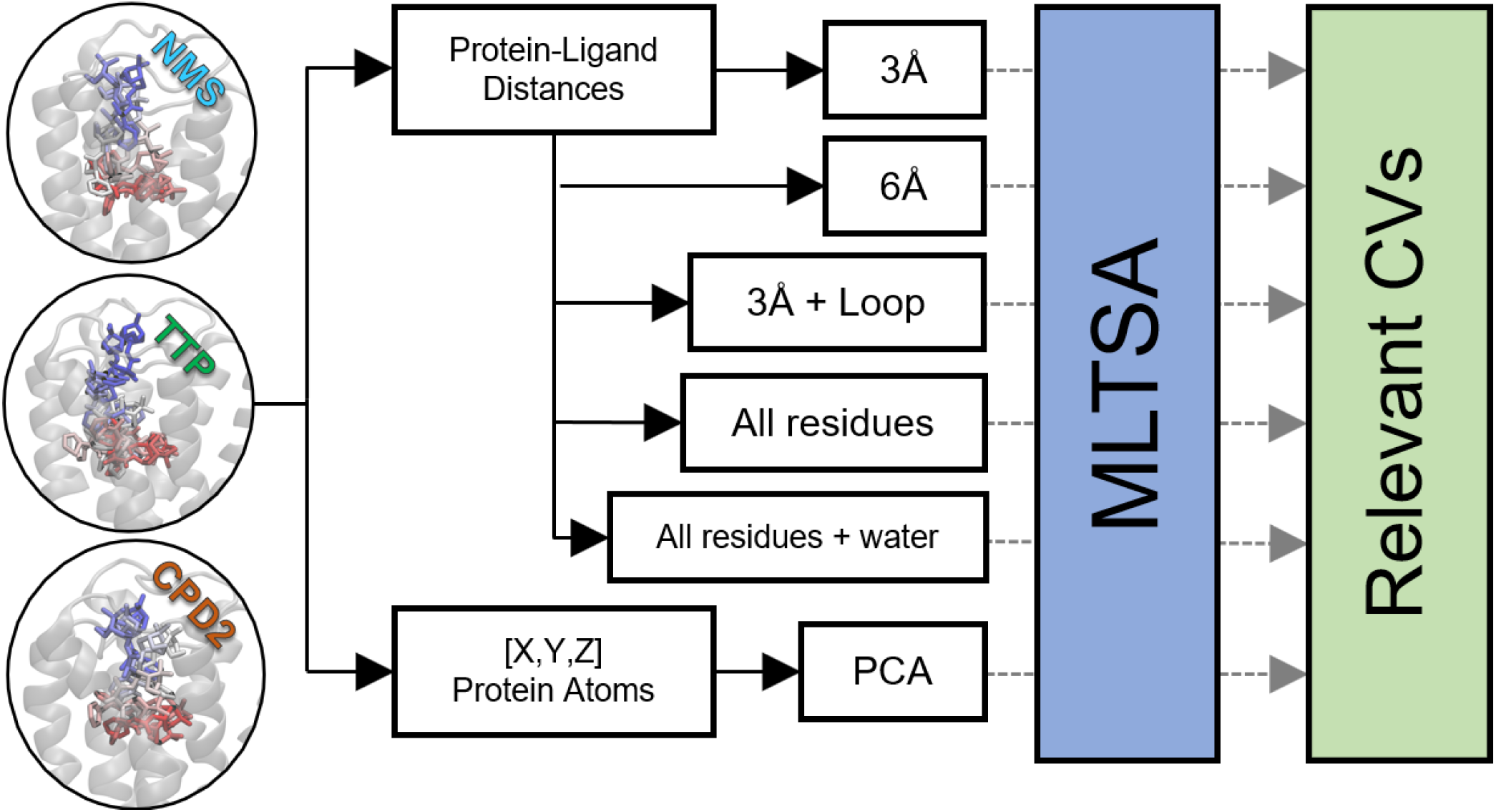

Graphical abstract of the work, showing the unbinding for ligands **1** (tiotropium, TTP), **2** (N-methylscopolamin, NMS) and **3** (homatropine methylbromide, CPD2). Using TTP’s downhill simulations from its unbinding transition state, different protein-ligand and proteinprotein interactions were analyzed with MLTSA to find relevant CVs driving the different outcomes.

## INTRODUCTION

Muscarinic receptors (MR) are a five-membered subtype group of transmembrane receptors, which form an important part of the parasympathetic nervous system. They are activated by neurotransmitters such as acetylcholine and muscarine,^1^ and transmit extracellular signals to the cell interior, which makes them attractive drug targets.^2^

The sequence identity between the five MR isoforms is low, except between the transmembrane regions.^3,4^ This region contains seven alpha helix substructures, which anchor the protein in the outer membrane of the cell.^5^ On the cytoplasmic side, the receptor is bound to a GTP-binding protein, which is responsible for the subsequent signal transduction. Therefore, MRs are part of the G-protein coupled receptor (GPCR) superfamily.

Downstream signaling can be spontaneously induced when MRs bind to GTP-binding protein, even in the absence of the corresponding agonist.^6^ Activation, as well as the downstream signaling can be suppressed when suitable antagonists are bound to MRs. This can be exploited pharmacologically,^7^ and several important muscarinic antagonists were developed and used for instance, as bronchodilators in the treatment of asthma or chronic obstructive pulmonary disease (COPD).^8–11^

Human MRs (hMRs) are expressed in a variety of tissue in the human body, therefore a drug with low selectivity may cause severe complications and side effects.^12^ While the hMR3 isoform - which controls the tension of the smooth muscle tissue in the bronchial tubes - is the actual target of bronchodilators, the off-target binding to the highly homologous transmembrane region of hMR2 is responsible for serious side effects, especially in the cardiovascular system.^12–15^ Due to the high homology between the two isoforms, the binding affinity of most muscarinic antagonists is very similar. For example, the pKi value of the pharmacologically widely used tiotropium for hMR2 is 10.7 and for hMR3 11.0.^12^ Nevertheless, tiotropium shows a high selectivity because the dissociation rate from hMR2 is significantly higher compared to that from hMR3 by about one order of magnitude.^12,16,17^ As a consequence, the residence time of tiotropium in the hMR3 isoform is very long and the binding was considered to be kinetically irreversible.^12,18^

As in general, the drug unbinding process is a rare event, it is highly challenging to study it experimentally and the detailed mechanism is still mostly unknown. However, there are several computational studies available that attempt to approach this problem via molecular dynamics (MD) simulations.^19,20^

Simulations on the beta-2 adrenergic receptor using RAMD found two different types of pathways for the unbinding of the beta blocker carazolol. One of them along the long axis directly into the extracellular space and one laterally into the membrane.^21^ Recently, it was shown that the path leading directly into the membrane is probably an artefact caused by the force constants of the biasing potentials being too high.^22^ For the same receptor, binding paths for several antagonists and agonists could be identified by conventional MD.^23^ A free energy profile (FEP) was also presented, which is characterized by two barriers. The first barrier describes the process of docking of the ligand from the solution to the tunnel entrance of the receptor (the extracellular vestibule). The second barrier is on the way of the ligand from the extracellular vestibule to the orthosteric binding site.

Later works using metadynamics and Markov State Models (MSMs) found the resting state in the extracellular vestibule to be very shallow and a significant barrier for the desolvation process could not be found.^24^ It is now largely consensus in the available literature that the rate determining step is indeed on the way from the vestibule to the binding site.^25,26^

Previous studies on the unbinding path of the hMR2 receptor and its agonist iperoxo have also shown that the process encompasses two steps. In these unbinding processes the rate limiting step was found to correspond to the ligand exiting from the orthosteric binding site to extracellular vestibule.^27,28^ Two different exiting pathways are suggested, the first one (and more favorable) involves the rotation of the ligand and its exit through the extracellular vestibule, while the second one is characterized by the rearrangement of the extracellular loop 2 (ECL2) limiting the ligand from entering the solvated state. Free energy profiles for the unbinding were estimated using metadynamics, however, calculations of the free energy barrier or unbinding rates proved to be challenging due to force field inaccuracies.^28^ Given the homology between hMR2 and hMR3 similar limitations are expected to arise, which have been considered for this study.

In this work, we applied our recently developed unbinding algorithm^29^ to hMR3, to investigate the dissociation of tiotropium (**1**) and two structurally similar ligands, N-methylscopolamin (**2**) and homatropine methylbromide (**3**) (Figure 1). The obtained unbinding pathways were refined using an adaptation of the finite temperature string method.^30^ Finally, the transition state (TS) of the tiotropium unbinding was detailed and analyzed with the aid of machine learning (ML) to identify prominent interaction pairs of the ligand and the receptor at different levels. Additionally, we also revealed key conformational changes of the protein that define the downhill trajectory outcomes.

**Figure 1.**
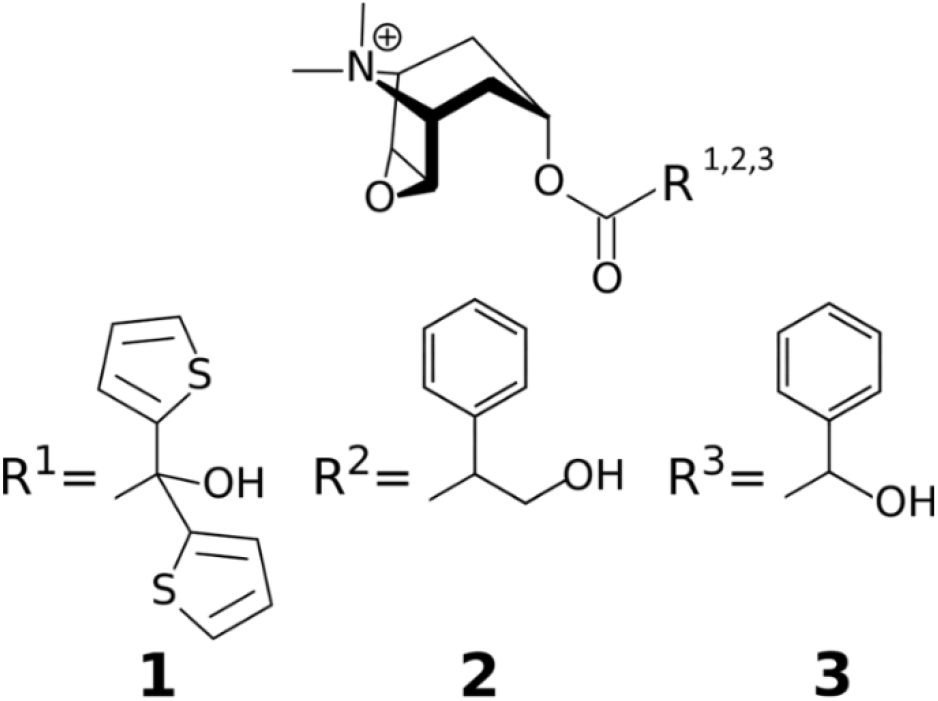
Structures of the ligands investigated in this study: tiotropium (1), N-methylscopolamin (2), and homatropine methylbromide (3).

## METHODOLOGY

### Starting structure

The starting coordinates for hMR3 were obtained using a rat MR3 crystallographic structure, PDB ID 4U14,^31^ with a resolution of 3.57 A and with tiotropium bound in the orthosteric site. Our structural model was truncated to the transmembrane helices and the extracellular loops, which are highly conserved between human and rat (91.85% homology), and contain the necessary and sufficient domains for ligand unbinding.^3^

### Parameterization and system building

The protein was inserted into a membrane using the membrane builder^32–34^ of the CHARMM-GUI web server.^35–37^ and then solvated in water^38^with 150 mM KCl. The membrane consists of POPC:DMPC:PYPE:DMPE in the ratio of 1:2:3:4, chosen on the basis of earlier studies of hMR3 and on tracheal membrane tissue.^39^

The ligands were geometry optimized at the B3LYP/6-31G** level of theory^40^ applying the ORCA 4.1 software suite.^41–43^ With the optimized structures, force field parameters for the ligand were defined using the CHARMM-GUI ligand reader.^44^

The all-atom CHARMM36m force field was used for the protein^45–48^ and the lipids,^49,50^ and the TIP3P model^38^ for the water. Simulations were carried out with the NAMD software package^51^ using input generated by the CHARMM-Input generator.^52^ The cutoff for non-bonded interaction was kept at 12 Å, the switch distance at 10 Å. Electrostatic interactions were handled by a particle-mesh Ewald solver with a grid spacing of 1 Å. The temperature was kept at 310.15 K using Langevin dynamics. Pressure was kept at 1.013 bar by Nosé-Hoover Langevin piston pressure control.^53,54^

The structures were first energy minimized according to the CHARMM-GUI scheme and subsequently equilibrated for 50 ns.

### Unbinding simulations

The unbinding procedure was followed as described in our previously published^29^ protocol. After the equilibration, a 20 ns production run without any restraints was performed. During this production run all interacting pairs of heavy atoms – one in the ligand and one in the protein – were identified. Thereby, a pair is defined as “interacting” if the distance between the atoms is below 3.5 Å for more than 50% of the simulation time. Based on the sum of these interacting distances, a collective variable (CV) is defined and restrained harmonically.^29^ During an iterative process, subsequent simulations of 10 ns use this biasing CV with a force constant of 10 kcal mol^−1^Å^−2^. The constraint position (i.e., the length) of the CV is monotonically increased. In the next iteration, new interaction sites are identified in the same way as before and these are added to the CV. Interactions are discarded and removed from the CV, if the distance between the atoms is larger than 11 Å. A shorter cutoff distance results in the ligand falling back into the original binding position after a few iterations. This procedure is repeated until the ligand is displaced out of the receptor.

The unbinding simulations were run for 25 iterations, adding up to a total of 240 ns simulation length. Thereby, a total of 52, 50, and 44 interacting protein-ligand distances were identified by our unbinding method along the paths for ligands **1**, **2**, and **3**, respectively.

### Refinement of the path using the string method

The unbinding path was used as a starting point for the following refinement using the finite temperature string method.^55^ Since the string iterations are computationally very expensive and at the same time converge rather slowly due to the many dimensions, only 20 iterations were calculated.

### Approximation of the TS region

To approximate a TS structure from the string windows, we identified a set of structures from the string windows, which are very similar in the unbinding paths of all three investigated ligands (Figure 2). We selected five windows as starting points around the window with these distinct structures for ligand **1** and performed 50 independent unbiased (downhill) MD simulations with 5 ns lengths each. Thereby, we were able to identify the structure that provided the closest 1:1 ratio of a binding (IN) or unbinding (OUT) events, which we considered to be the TS of the unbinding process.

**Figure 2.**
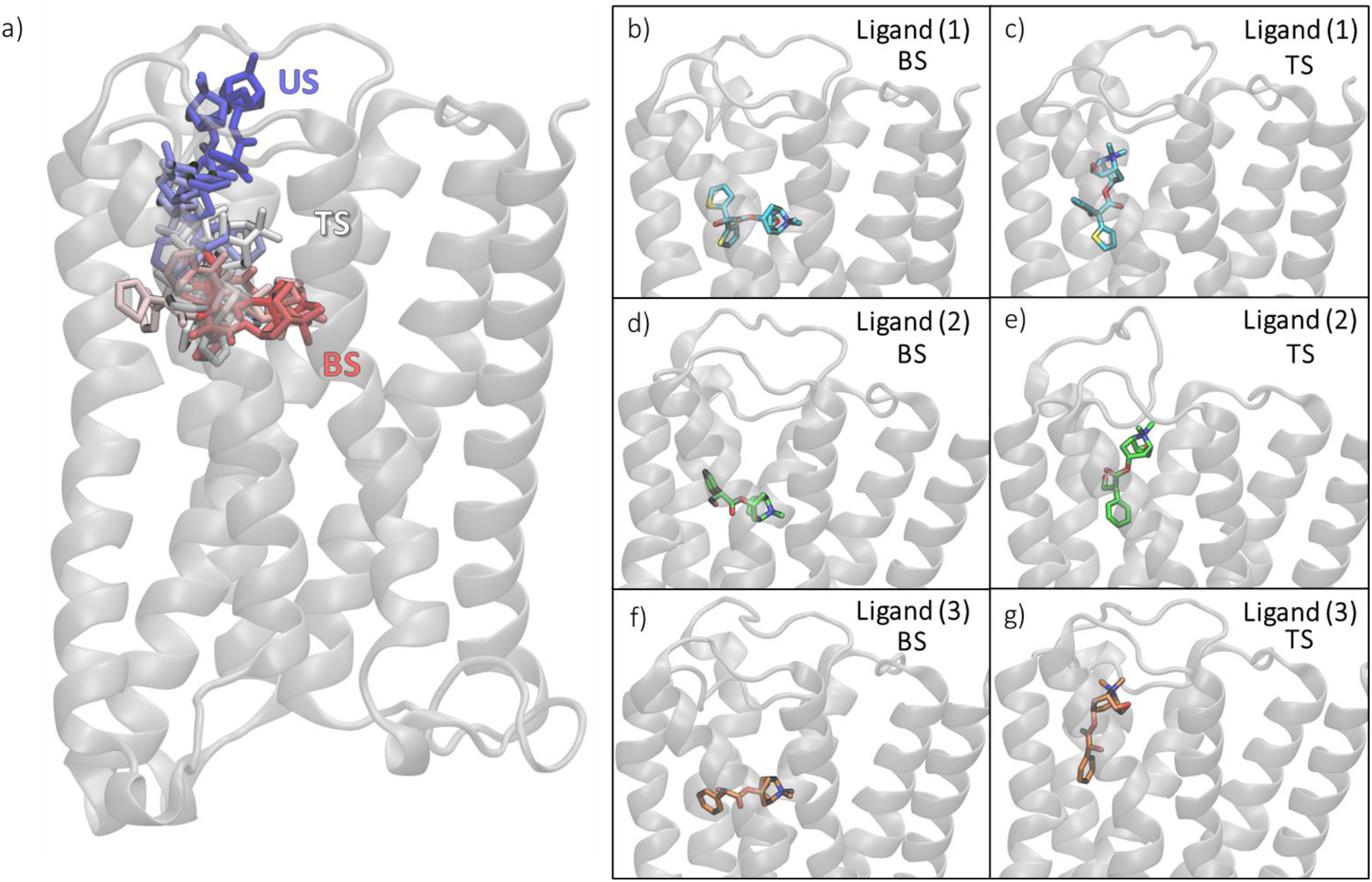
Left: a) Overlay of the structures from the unbinding path of ligand 1 through time starting from the bound state (BS, in red) towards unbound state (US, in blue) passing through an approximated transition state (TS, in white). Right: stick representations of the three unbound ligands on their original BS (b, d and f, for ligands 1,2 and 3 respectively) and their TS (c, e andg, for ligands 1,2 and 3).

### Machine Learning Transition State Analysis (MLTSA)

To aid the identification of the main CVs driving the system across the TS and to pinpoint novel descriptors that determine the fate of a binding/unbinding events, we used our MLTSA^29^. In this approach, we train an ML model to predict the outcome of downhill simulations with data close to the TS. Subsequently, we make use of the trained models to discover the key TS-defining features of the system.

### Creation of the datasets

Using ligand **1**’s identified TS structure as the starting point, we ran multiple 5 ns long unbiased simulations. We classified and labelled 149 downhill trajectories by considering a linear combination of 52 distances to identify which simulations arrive at an IN or an OUT state. A minority of additional trajectories not reaching clearly either the IN or the OUT states after 5 ns were discarded. To train ML models, we created several sets of features containing different distances (CVs) along the simulation frames. To assess intra-protein interactions, a first dataset (*XYZ-PCA set*) included the Cartesian coordinates of all protein atoms (~6K, not including hydrogens). To reduce the dimensionality, we applied principal component analysis (PCA) and used only the top 100 components as features. To enable more interpretable localized features, we created further datasets containing ligand-protein distances. The first such set (*3Å set*) contained all interatomic distances between the ligand and the protein within 3 A of the ligand at the starting TS position, excluding hydrogens. The second dataset of this type (*6Å set*) was created in a similar fashion to the previous one, but with a cutoff of 6 A instead. For the third dataset (*3Å+ECL2/TM5 set*), the same data was used within 3 A of the ligand, with the addition of the interatomic ligand-protein distances of the extracellular loop 2 (ECL2) and the transmembrane region 5 (TM5), including residues from I222 to T231. An additional dataset, to assess overall ligand-protein contributions, was also created (*allres set*), which considers all residues and includes the closest distance between the residue and the ligand at each simulation frame. This dataset was also amended with the closest 8 water molecules, their distances to the ligand (*allres+wat set*) was included to enable the assessment of the role of water molecules.

### Machine Learning models and training

We used two different ML models: a Multi-Layer Perceptron (MLP) neural network classifier,^56^ and a Gradient Boosting Decision Tree (GBDT) classifier.^57^ Both models were trained to predict the outcome (IN/OUT) of the simulations from early on data at the time range from 0.05 ns to 0.1 ns, totaling 2500 frames per simulation. We trained 100 independent MLP and GBDT models randomly assigning the 149 simulations into training data (70%) and validation data (30%). Details on the trainings and hyperparameters can be found at SI section 2.

### Feature Analysis

We used the Gini feature importance^58^ to evaluate the relevance of the features from the GBDT models, averaged across the 100 trainings to calculate their relative feature importance (RFI). To identify key features in MLP models, we removed the variance from each feature one-by-one^29^ and assessed the accuracy drop they encounter when predicting outcomes with the trained models. If the accuracy of the prediction is greatly reduced when a feature is altered, the feature was considered important for the description of the TS. We identified the overall top features averaging the relative accuracy drop (RAD) from all 100 trainings on all datasets used (Figures 3–5).

**Figure 3.**
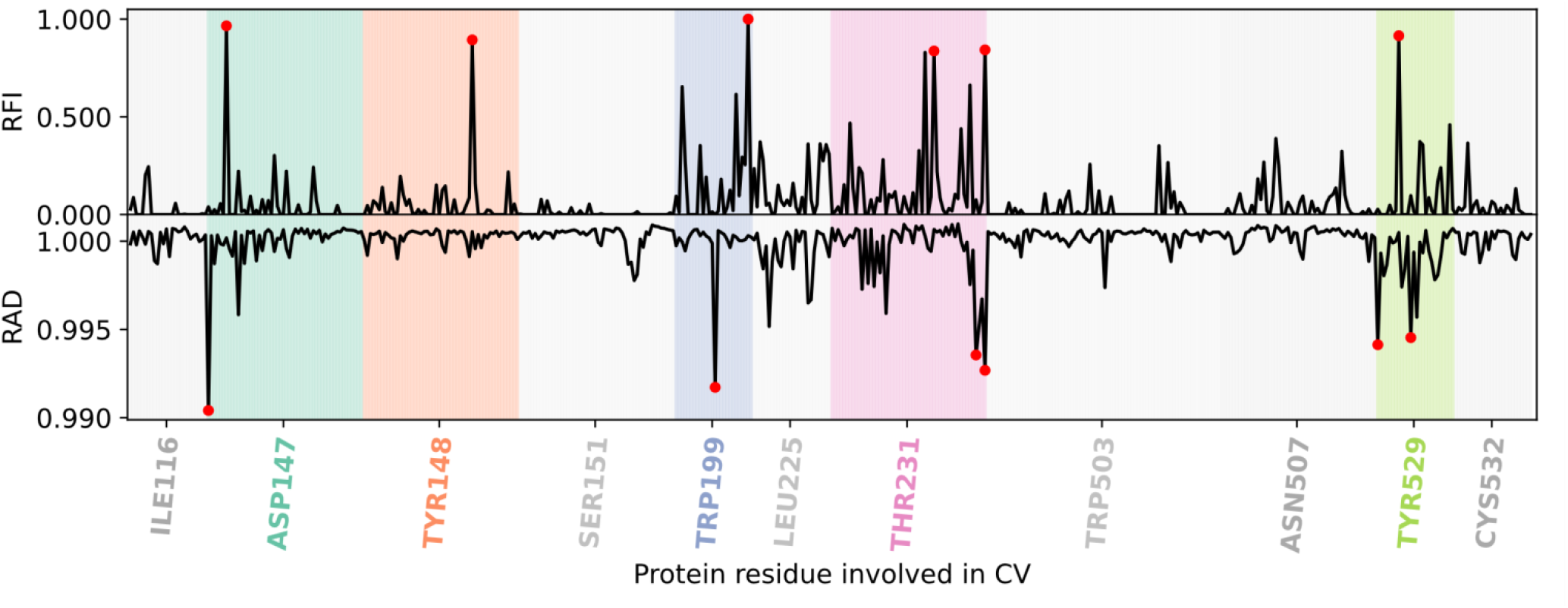
Relative feature importance (RFI, top) and relative accuracy drop (RAD, bottom) shown for every interatomic distance between ligand 1 and hMR3 in the 3Å dataset. Distances are ordered and clustered by residue number. Residues with the top six distances (redsymbols) are highlighted.

**Figure 4.**
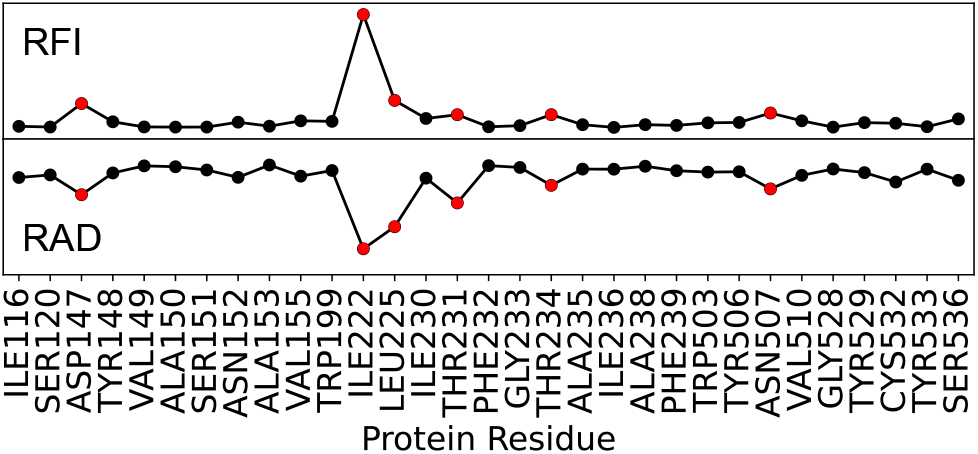
Average RAD (from MLP) and RFI (from GBDT) of the interatomic distances of the ligand 1 per protein residue for the 6Å dataset. In red, the top 6 residues detected by both approaches.

**Figure 5.**
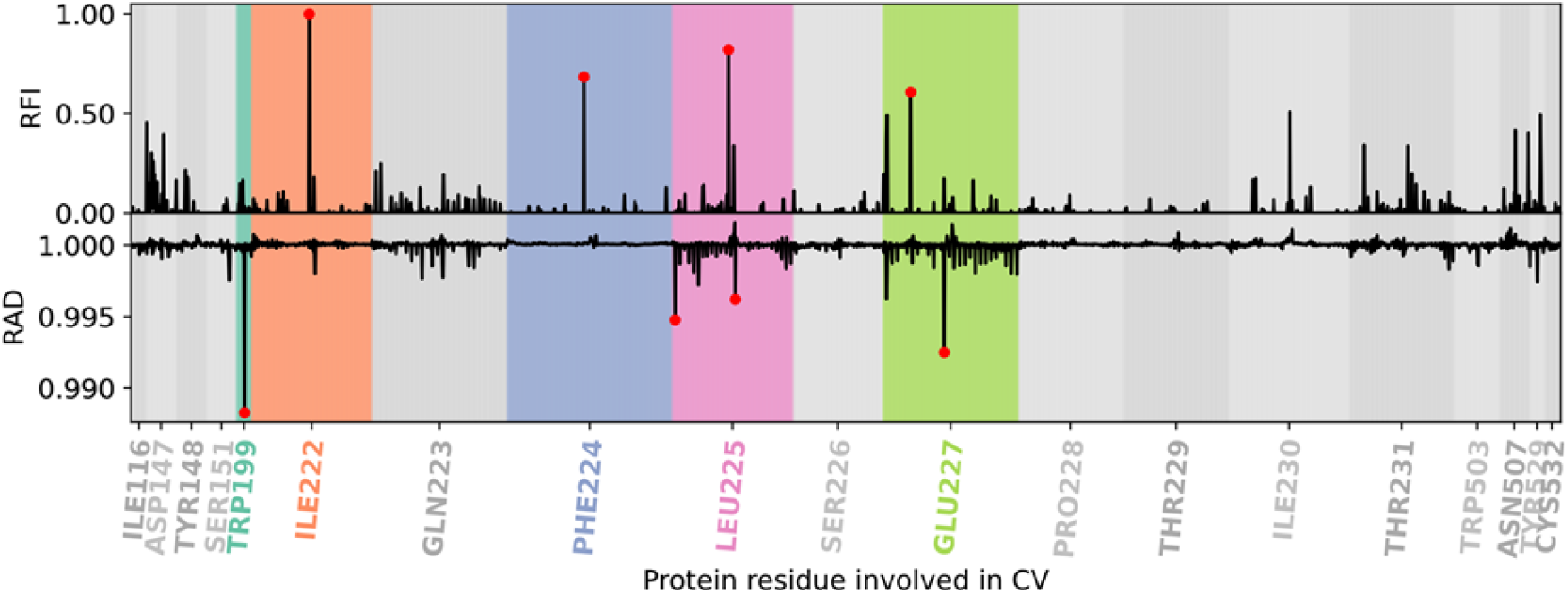
Relative feature importance (RFI) (from GBDT model) and relative accuracy drop (RAD) (from MLP model) values for each interatomic ligand-protein distance per residue in the ligand 1’s 3Å+ECL2/TM5 dataset. Marked in red are the top distances for each model. Highlighted, the most important residues for the ML models.

## RESULTS AND DISCUSSION

### Bound state in the orthosteric site

In all three ligands, the initial ligand positions in the unbinding simulations are close to the starting bound pose: the charged end of the molecule is nestled in an aromatic cavity, which is formed by the residues W503, Y148, Y506, and Y529. The tyrosines form a cap around the ligand. Simultaneously, the S151 residue coordinates the epoxide group via a hydrogen bond and the negatively charged residue D147 neutralizes the positive charge of the ligand. At the opposite end of the molecule, the N507 residue stabilizes the molecule by a hydrogen bond with the OH group. The same binding mode was also described in recent works.^18,25,31,59^

### Departing from the binding site

As illustrated in Figure 2 and S1, the first movement from the binding state (Figure S1.**A**) is a rotation of the charged end of the molecule. Thereby, the hydrogen bond of the epoxide group with S151 is broken and the ligand slightly gains flexibility. Apart from that, the ligand’s position in the binding site remains nearly unchanged (Figure S1.**B**). This first movement is most pronounced for ligand **1**, which follows a helical motion along its longitudinal axis and thus it detaches itself from the aromatic cavity.

This shift is present, but less pronounced for ligand **2** (Figure S2a). Subsequently, ligand **2** breaks through the tyrosine-formed ceiling via a path associated with significantly more dislocation of the residues Y148, Y506, and Y529.

In the path of ligand **3**, the entire molecule does not shift, instead mainly the end of the ligand with the thiophene ring moves (Figure S2b). This allows the charged end to slip outwards the aromatic cage in a rolling motion.

### Through the bottleneck

The new position after the shift, allows the molecules to rotate their charged end by 90° towards the direction of the receptor tunnel’s exit (counterclockwise), without exerting a lot of tension on the tyrosine residues forming the aromatic cap. During this rotation, all three paths pass through a state (Figure S1.**D**), which is highly similar in all unbinding trajectories. Interestingly, this rotation was observed to proceed clockwise for the iperoxo ligand unbinding path in hMR2.^28^ This movement positioned the charged end of the molecule pointing towards the membrane in these previous simulations, and not to the extracellular vestibule observed by us. We subsequently found by unbiased simulations starting from this structure that either the ligand is led back into the binding site, or it moves along the exit tunnel towards the extracellular vestibule (Figure S1.**E**). Therefore, this position can be identified as a TS of the unbinding from the orthosteric site. The pathway is also similar to the previously reported forced dissociation with acetylcholine as well as tiotropium (**1**) on hMR3, and with a slightly tilted orientation on hMR2.^25^

### To the extracellular vestibule

For all ligands the total length of the simulations was not sufficiently long to observe the complete unbinding of the ligand, rather the ligand remains in the extracellular vestibule but outside the orthosteric binding site. However, in line with the consensus literature, it is estimated that the final unbinding step from the extracellular vestibule has a significantly lower barrier, therefore it does not likely contribute to the off rate.^24,25^

### Downhill trajectories from TS structures

We evaluated starting structures from 5 string windows near the bottleneck conformations. The structure closest to the TS position led to 85 and 64 downhill trajectories of 5 ns reaching the IN and OUT states, respectively (Figure S3). To explore the time range where the TS is probed, we performed initial ML trainings to identify the region where the ML method can accurately, but not with full confidence, predict the final outcomes from as early timeframes as possible. We found that this was already possible from 0.05 to 01 ns timeframes. Trainings at different times can be found in Figure S5, final accuracies for all datasets in Table S1.

### Assessing contributions from protein conformational changes

To consider changes in the protein structure affecting the unbinding, we analyzed the protein Cartesian coordinates via their top 100 PCA components (Figure S4). We were able to predict the outcome very accurately, obtaining average test accuracies of 100% (MLP) and 93% (GBDT). Out of the 100 components, the first two PCA components were important both for RFI and for RAD. Additionally, PCA23 and PCA59 were important for RFI (see SI section 2). The main PCA component represents large-scale movements from the TM2, TM3, and TM6 to TM7 helices, including some ECL1 residues (Figures S6 and S7). The residues that contributed the most are from the middle of TM6, close to the ligand. The second main PCA component (top RAD feature) represents motions from the rest of the protein, mostly from TM4 to TM5, with the ECL2 loop being especially relevant. The largest contributions come from residues (W206, Q207, I222 and Q223) that belong to ECL2/TM5 junction, some from TM4 that are close to the ligand (I194 and V193). However, due to the broad distribution present in the PCA components, their interpretability is limited. Therefore, we next focused on feature sets that are precisely localized and able to assess specific ligand – protein atomic distances instead.

### Key feature identification from the 3Å dataset

We created a high-resolution dataset, which contained atomic distances between the ligand and protein residues within 3 Å of the TS structure of the ligand. Using our 3Å dataset, we achieved a prediction accuracy of ~78% with MLP and ~77% with GBDT, and obtained consistently similar key features by RAD and RFI (Figure 3). Both models (MLP and GBDT) agreed on the importance of four out of six top residues: D147, W199, T231 and Y529.

Three of these key residues were previously known to play important roles in the unbinding process. D147, as mentioned earlier, interacts with the charged amine moiety in the bound form. Similarly, Y529 is part of the aromatic cage around the ligand. Additionally, the aromatic substructures of the ligand are known to interact with a hydrophobic region close to W199. Mutational studies show an accelerated dissociation for Y529A and reduced half-life for both W199A and D147A, further suggesting their involvement. ^18^

Interestingly, T231 was not previously reported and validated as relevant for ligand interactions in the bound state.^18^ Even though there are no experimental studies, it was previously identified computationally to form relevant contacts during the forced dissociation of tiotropium.^25^

### Contribution of the extracellular vestibule within 6 Å

To assess the contributions from more distant atoms beyond 3 Å, we also analyzed results from a dataset that includes ~5000 interatomic distances within a range of 6 A from the ligand at the TS. In this dataset, we analyzed both individual feature importances (Figure S8), and average importance values for each residue (Figure 4). Accordingly, D147 and T231 are again part of the top 6 key residues both measured by RAD or RFI. Newly identified key distances include I222, and T234, which were not part of the previous dataset, as well as additional heavy atom distances from L225 and N507. L225 was previously reported relevant for the binding/unbinding kinetics in hMR2/hMR3 experimental studies, but insufficient alone to explain the difference between both receptors. N507 is a previously validated relevant interaction that accelerated the dissociation of tiotropium when mutated to Ala (N507A).^18,25^

Interestingly, the residue with the most relevant interactions is I222 and it was not described previously. Together with L225 and T231, I222 forms a hydrophobic cluster on the extracellular vestibule (Figure 6 and 7a, b, i). The fact that the most relevant residues (I222, L225, T231, T234) are close together in an extra-cellular loop (ECL2) may be indicative of the importance of this loop for the unbinding. When aligning hMR2 and hMR3 protein sequences, most of the sequence is identical, but the region prior to T231 (ECL2/TM5) has a high genetic variability (Figure 8). Interestingly, preceding I222, there is another variation in the sequence for ECL2: F221 in hMR3 is substituted with Y177 in hMR2. Moreover, this residue is a potential phosphorylation/modulation site for hMR2,^60,61^ thus thought to be not only an important region for allosteric regulation, but it could alter the observed unbinding kinetics depending on the phosphorylation state of hMR2. This suggests that the residues between I222 and T231 may be relevant to the significantly different behavior observed between hMR2 and hMR3 in terms of residence times.^12^ Hence a third dataset (3Å+ECL2/TM5 Loop) was created containing all the residues prior to T231, which range from I222 to T231.

**Figure 6.**
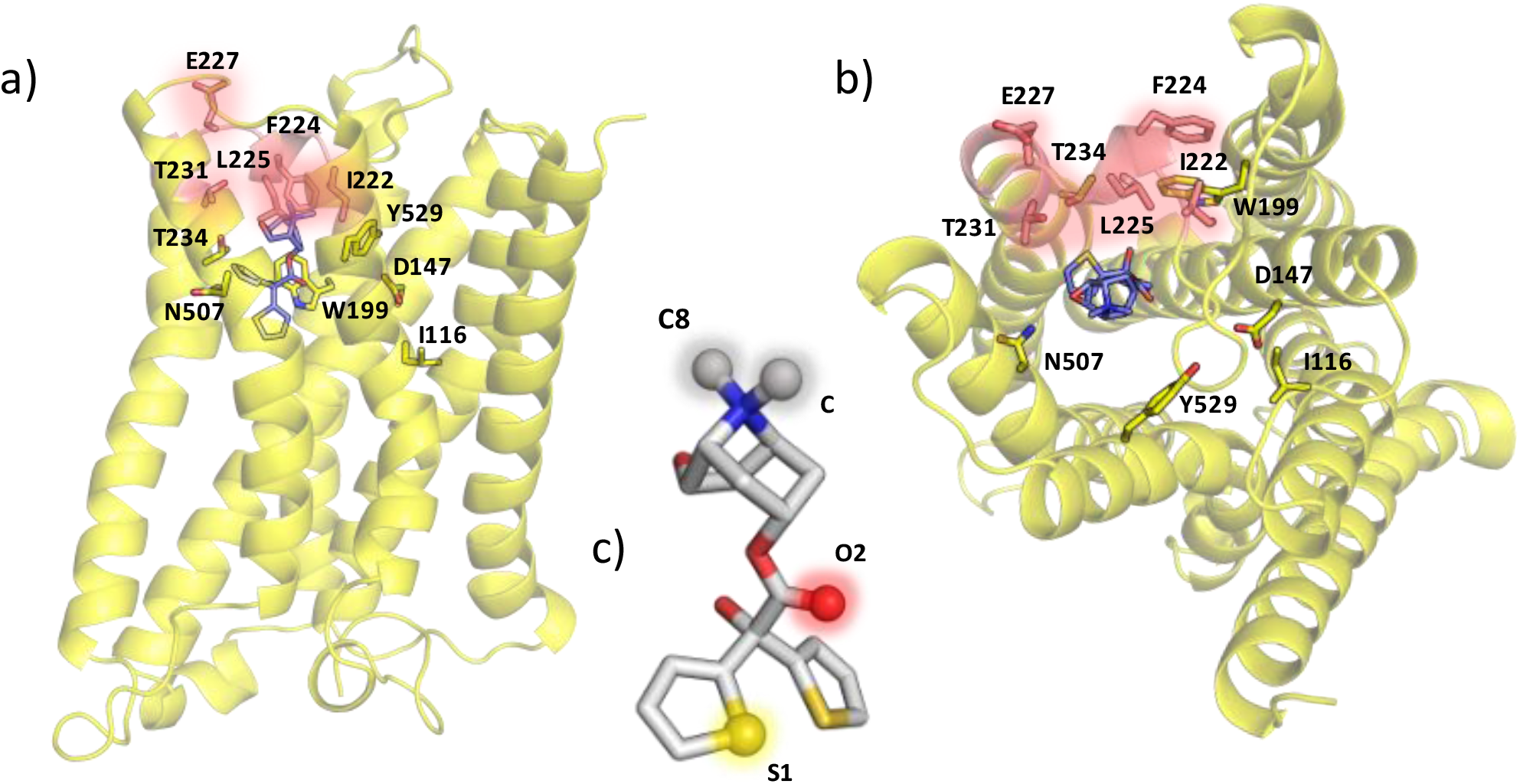
Top: Front (a) and top (b) view of the M3 receptor at the TS, in sticks the most relevant residues for the unbinding process found by the ML models. In salmon, the residues belonging to the ECL2 loop, which is found to be the most relevant region. c) Ligand 1’s structural representation with the most relevant atoms found by the MLTSA, highlighted, and annotated.

**Figure 7.**
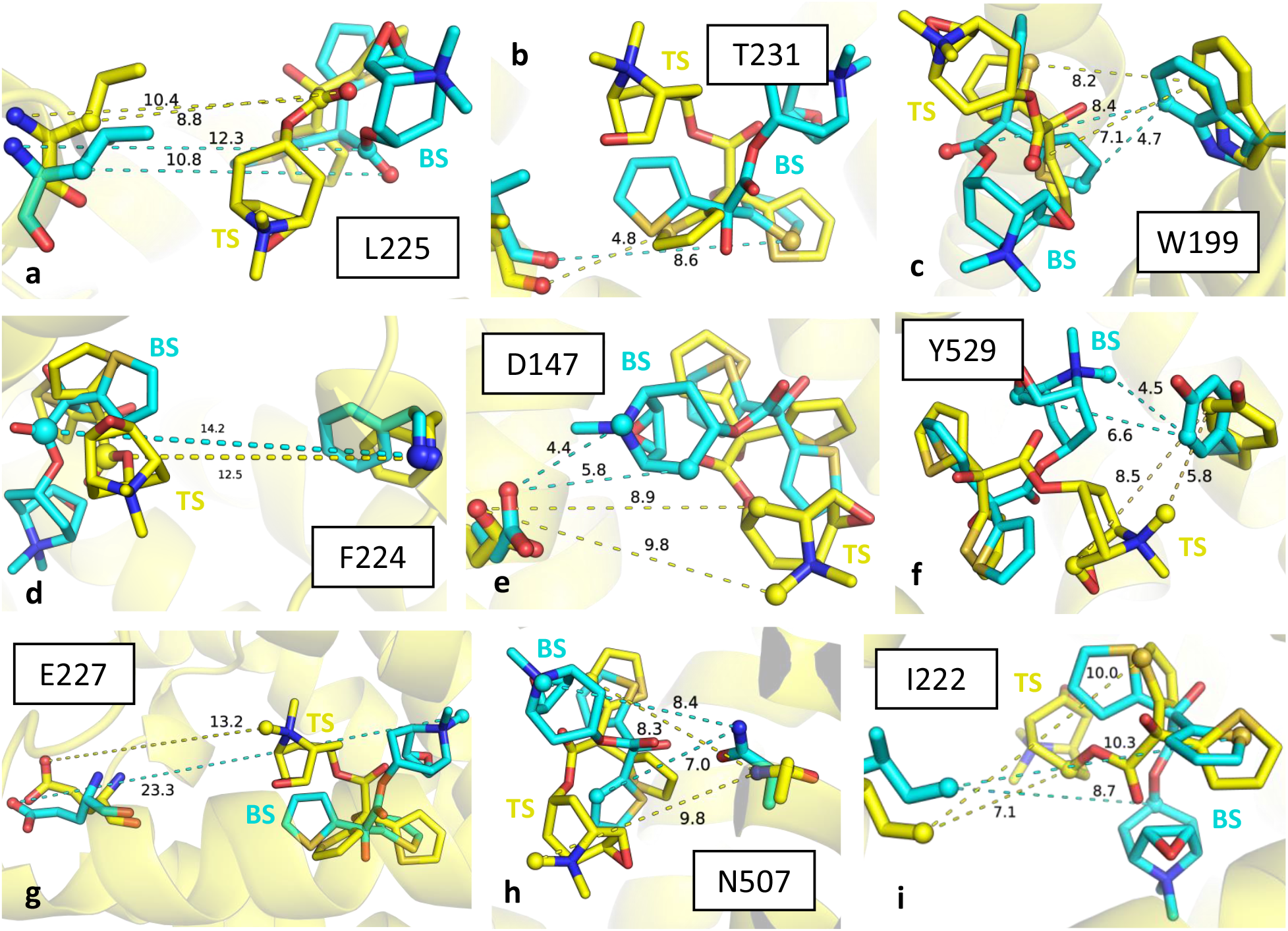
a) to i) are the top nine residues represented as sticks with their protein-ligand (hMR3-ligand 1) distances consistently found to be most important throughout the MLTSA analysis across all datasets. In yellow, the ligand-protein complex at the TS and their distances, in cyan the ones corresponding to the complex at the BS. Represented as spheres, the atoms that the interatomic distances represented correspond to.

**Figure 8.**
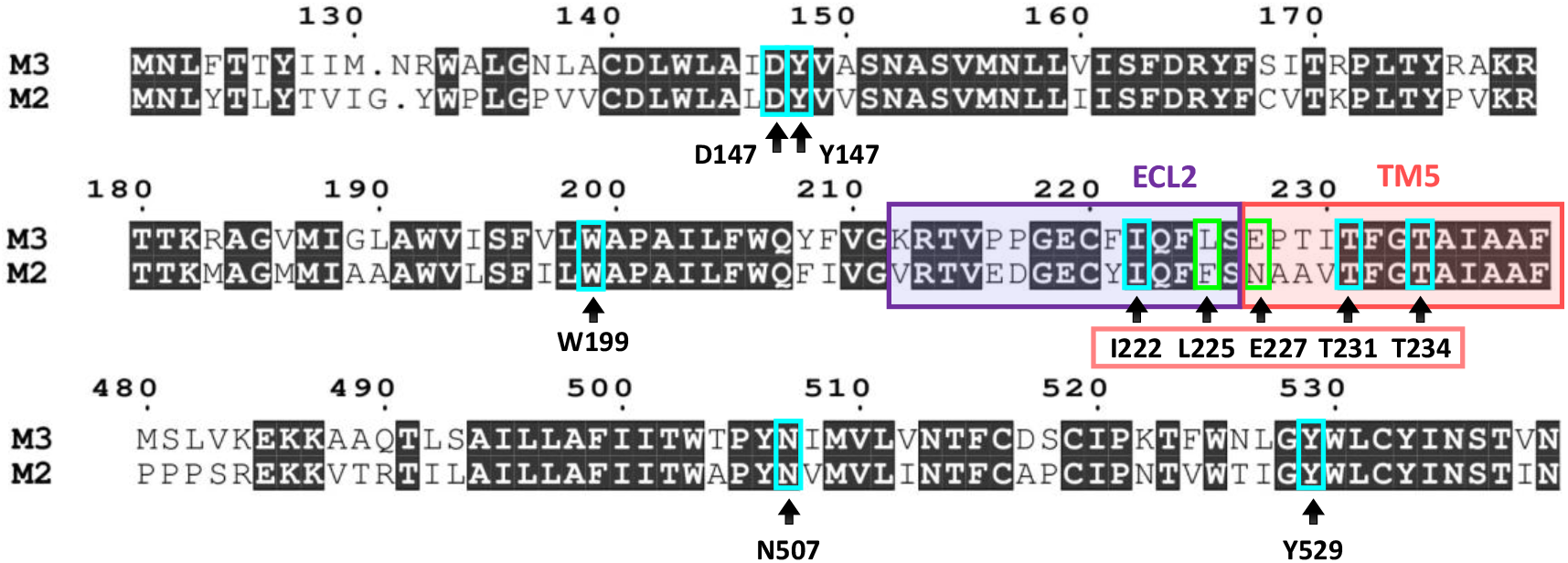
Protein sequence alignment of hMR2 and hMR3 for selected regions involved in the unbinding process. Key residues identified by MLTSA are distinguished as conserved (red) or non-conserved (green) between the two receptors. The ECL2/TM5 region is also highlighted (purple and salmon).

### Exploring the role of the ECL2/TM5 junction

In the presence of distances from this region (Figure 5), the top features belong mostly to the ECL2/TM5 junction, except for W199 when using RAD. In hMR2, L225 corresponds to a Phe residue (Figure 8), which is bulkier. Interestingly, this change was previously reported to remove a pocket in hMR2, which is present in hMR3.^25^ The negatively charged E227 is replaced by a neutral Asn in hMR2. Remarkably, both ML models found E227 important, despite its longer distance (Figure 7g). This residue has been mutated to Ala (E227A) previously, resulting in a slight decrease in the half-life of tiotropium, **1,** from 24.5 h to 20.1 h. The RFI, however, found an additional key distance involving F224 as one of the most relevant distances. When mutated to F224A, the half-life of **1** is reduced by ~50% to 13.8h.^18^

Additional tests with distances from an alternative loop, ECL3, were also added to the 3A dataset and analyzed (Figure S9) for comparison. These demonstrate no significant contributions from this region, thus validating the unique role of the ECL2/TM5 junction.

### Structural spotlights of tiotropium involved in unbinding

Our results point to key atomic contributions from only a few selected atoms of tiotropium (Figure 6c). The most prominent moiety corresponds to the methyl groups (C and C8 atoms) that are bonded to the charged amine. Of key relevance is also the S1 sulfur atom from only one of the two thiophene rings, showing key interactions with W199, I222 and T231 (Figure 7 panels c, b and i, respectively). Finally, the O2 atom from the carbonyl oxygen of the ester group is also important, as identified in interactions with W199, L225 and Y529 (Figure 7 panels c, a and f, respectively). In agreement with our results, previous studies have shown that the tiotropium analogues with the closest Ki values have a pattern containing all three groups: an amine cap, the carbonyl group in between, and two aromatic rings (thiophene or not) at the end.^18^

### Overall residue-ligand contributions

To assess all the residues in the protein, we decreased the resolution of the feature space, and evaluated only features defined via the closest distances between each residue and the ligand (*allres* dataset). This allows us to evaluate all residues, including the ones far from the ligand, which can nevertheless have key impact on the simulation outcome. The resulting training from this dataset yielded ~79% for GBDT and ~77% for MLP on their test set. T234, highlighted in our previous results as a key residue in the 6A set as well, is the most important feature for RAD, and second most important for RFI, validating its key role (Figure 9a).

**Figure 9.**
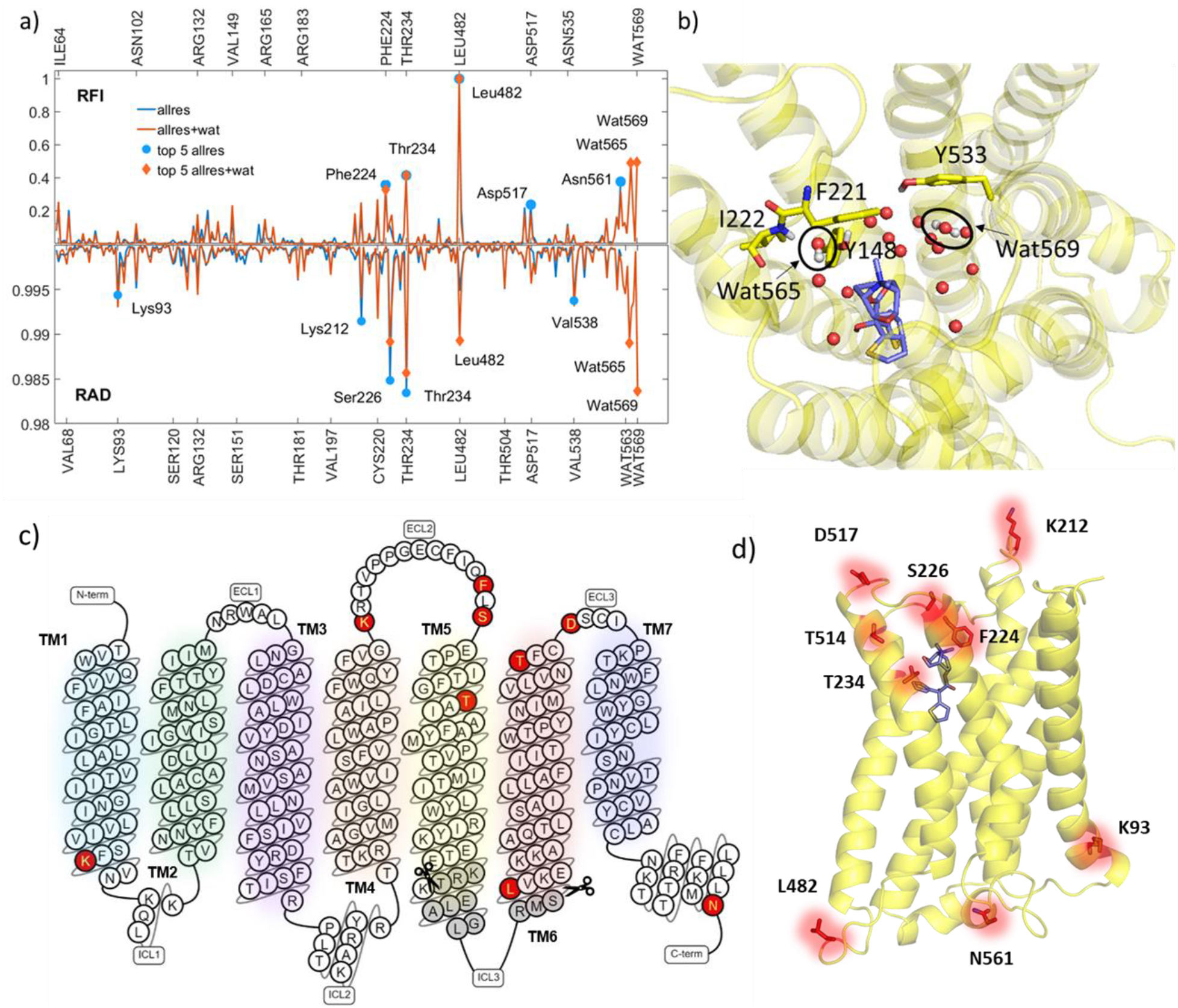
a) RFI and RAD for the allres (blue) and allres+wat (orange) datasets, highlighted are the top 5 residues for each approach (blue circle and orange diamond, respectively). b) TS snapshot showing the top two water molecules as well as nearby residues as stick in the allres+wat dataset. c) Diagram representation of the sequence of hMR3 portraying the different secondary structure motifs. In red, the top residues found decisive for the outcome by our MLTSA. In grey, the residues (kinase domain) not included in our simulation system. d) Top important residues from MLTSA highlighted in the 3D representation of hMR3, mostly corresponding to the ECL2/TM5 junction and the different ends of the alpha helices throughout the receptor.

A more distant residue that shows key importance is L482, ranked 1^st^ for RFI and 5^th^ for RAD. This distant residue is at the N-terminal end of the TM6, located very near the kinase domain of ICL3, at the interface of membrane and the intracellular matrix (Figure 9c-d). This could signal changes in the ligand-bound state to the ICL3, which is not modelled in our simulation system. Accordingly, this region is located between two main binding regions of hMR3 for activation and regulation.^62,63^ Pyrophosphatase-2 (PPase 2A), a transmembrane enzyme which targets the C-terminal region of the ICL3, the “**KRKR**” motif in (“IT**KRKR**MSLIKEKKAAQ”), is thought to be involved in hMR3 dephosphorylation.^64^ Additionally, the muscarinic receptor signaling regulator, SET, a PPase 2A inhibitor, also binds to the same motif.^65^ Furthermore, it was also suggested that protein kinase G II (PKG-II) activates hMR3 via a cGMP-dependent phosphorylation at S481 (“MSLIKEKK” motif).^62,66^ Therefore this region is thought to be a putative phosphorylation site just preceding L482.^62^ Interestingly, ligand-dependent phosphorylation of S481 was also connected to enhanced dimerization and/or oligomerization.^67^ This has been suggested previously in conjunction with homologous GPCRs,^68–70^ pointing to a general signaling mechanism in this family of proteins.^71,72^ Homo or hetero dimerization of kinase domains is often observed functional requirement along with phosphorylation when activating signaling pathways in general.^73,74^ L482, however, is the first residue in our simulation model after the missing kinase domain, hence the precise role of signal transduction from the orthosteric site to the ICL3 kinase domain remains to be explored in more detail.

Other key residues also include distant locations that are near the ends of helical domains, similarly to L482: K93, K212, T514, D517, and N561. Some of these residues were identified as important by mutational studies, such as K212V and D517A, that decrease the tiotropium residence time in hMR3. Near T514 and D517, C519 was also previously identified as a key residue for RFI in PCA components 23 and 59.

The ECL2 loop remains key in this dataset as well, besides T234, F224 and S226 are also highlighted (Figure 9a). This is validated by the F224A construct, as mentioned earlier, where the half-life of **1** is halved.^18^ In summary, RAD and RFI show a consistent picture, pointing to the key relevance of the ECL2/TM5 junction, in agreement with our previous results.

### Water plays a role in the unbinding path

Solvent molecules are known to play a crucial role in ligand unbinding kinetics.^75–79^ By both enabling the favorable electrostatic environment and orchestrating movements via hydrogen bonding, water molecules play a role that is often difficult to elucidate. To explore the role of water during the unbinding process, we included the 8 closest water-ligand distances together with the *allres* dataset as additional features. We found a modest increase in both MLP and GBDT prediction accuracy (~81% and ~79%, respectively). With these additional set of features, both RAD and RFI ranked the same two water molecules (Figure 9a-b, labelled 565 and 569) within the top 5 features. L482 remains ranked 1^st^ for RFI, and it is 5^th^ for RAD. Both approaches consistently find L482, T234, F224 and S226 most relevant together with water molecules (Figure 9c-d). This finding suggests that movements of water molecules in the pocket are also decisive to ligand unbinding in addition to the residues highlighted previously.

Water 565 is located near the ECL2 residues F221, I222, and forms H-bonds with Y148 and the backbone of I222, part of the ECL2/TM5 junction we highlighted throughout this work. Upon analyzing the most likely distances for IN and OUT trajectories, we observed that this water gets displaced in most of the trajectories as the ligand enters the orthosteric site. On the other hand, water 569 is on the other side of the ligand, closer to Tyr533, as well as Tyr529, which also forms the tyrosine cage. It only partially forms H-bonds with other water molecules, and it is located near a hydrophobic region of the pocket. While for OUT trajectories this position is not likely to change significantly, for IN trajectories the water moves deeper into the binding pocket as the ligand moves down into the orthosteric site.

## CONCLUSIONS

We generated and obtained consistent unbinding paths from hM3R for three ligands: tiotropium (**1**) and its analogues, **2** and **3**. All three ligands showed similar unbinding paths, including a first rotation of the charged end and a movement of the aromatic rings of the ligand, followed by a dislocation of the tyrosines forming the aromatic cage, finishing with a 90° angle rotation corresponding to the bottleneck while moving towards the receptor tunnel. Therefore, all ligands show a well-defined similar TS position and leave the orthosteric site in a highly homologous mechanism. The main barrier contribution in the unbinding process is known to be related to the ligand leaving the orthosteric site,^18,23,25^ therefore we did not follow up the subsequent full exit out of the vestibule. Our obtained paths are also in agreement with previous studies on the unbinding paths of tiotropium for both hMR3 and hMR2, where the ligand exits in a similar way.^18,25^

We further validated our TS structures by generating unbiased downhill simulations, which allowed us to further analyze the main events driving the unbinding at the TS. Our first Cartesian coordinate-based (XYZ-PCA) dataset showed a remarkably good accuracy at predicting the outcome of the simulation at very early times. This first analysis suggested relevance of the ECL2 loop and the residues at the ends of the transmembrane helices but proved hard to interpret. A more local but high-resolution (3A) dataset, which included the relevant protein binding pocket – ligand atomic distances at the TS structure, matched experimentally relevant residues such as D147, Y148, Y529 and pointed to T231 which is part of the ECL2/TM5 junction. An increased dataset (6A) continued to point towards ECL2/TM5 junction contributions being the most relevant. We further tested the relevance of this region by augmenting our previous 3A dataset with these residues (3A+ECL2/TM5). This further justified the key role of the ECL2/TM5 junction. On the other hand, adding e.g., ECL3 residues to the 3A dataset instead did not yield relevant distances from the ECL3 region. This further validated the relevance of the highlighted residues from ECL2/TM5, which also show differences in the protein sequence compared with hMR2 (L225/F181 and E227/N192 substitutions), highlighting potential role in the residence time differences between the two receptors.

Several residues identified by the MLTSA were previously experimentally mutated, further validating their importance in residence time. The available mutations show the largest influence for F224A, Y529A and N507A in the unbinding kinetics, while D147A, W199A, E227A, K212V and D517A impact it to a lesser extent. Additional residues we identified here as highly relevant remain yet to be experimentally probed for their role in ligand unbinding kinetics, such as L482, together with the preceding S481, as well as T234 remain to be further studied. Other identified residues that could play a role are: C220, I222, L225, S226, T231.

Our results point to the structural importance of key ligand groups and consistently found specific atoms in the amine end, the carboxyl group, and the tiophene rings, to be highly relevant. All three pharmacophore groups match other variants of tiotropium that have a charged end, middle carboxyl group and an aromatic ring at the end, either one or two.^18^ Our analysis can therefore provide useful information to propose pharmacophores in future drug design studies for kinetics-based ligand optimization.

To account for all residue interactions with the ligand, a dataset with coarser interaction features (*allres*) was also used. This confirmed the importance of the ECL2/TM5 junction, and furthermore pointed to residues at helical ends. Additionally, when the closest ligand-water distances are added to the previous set (*allres+wat* set), two water molecules also appear at the top. Our results suggest an important role of these molecules, whereby their movement is highly correlated to the ligand entering the orthosteric binding pocket. Importantly, L482 remains to be a top-ranked feature, near a phosphorylation site (S481 for PKG-II)^62,66^ and between two specific binding regions for signaling and activating proteins (SET and PPase 2).^64,65^ Interestingly, S481 phosphorylation was linked to enhanced dimerization in an allosteric mechanism upon antagonist binding,^67^ proposed to be a general mechanism in GCPR signal transduction.^68–70^ This suggests that the conformational changes of the ECL2/TM5 junction at the TS crossing transduce a signal across the membrane to the intracellular ICL3 kinase domain of the receptor as the ligand exits or binds the orthosteric site. Our MLTSA analysis appears to capture and identify allosteric effects, opening up potential avenues in various other systems and processes as well,^80,81^ beyond ligand unbinding. Nevertheless, the allosteric signal transduction remains to be studied in more detail, to aid the understanding of the function and mechanism of this biomedically-relevant receptor family.

## Supporting information

Supplementary Information with Tables/Figures

## ASSOCIATED CONTENT

### Supporting Information

The Supporting Information is available free of charge on the ACS Publications website.

## AUTHOR INFORMATION

### Author Contributions

The manuscript was written through contributions of all authors. / All authors have given approval to the final version of the manuscript.

### Funding Sources

We acknowledge funding by EPSRC (grant no. EP/R013012/1) and ERC (project 757850 BioNet).

### Notes

The authors declare no competing financial interest.

## ACKNOWLEDGMENT

We thank D. Callum for helpful comments on our manuscript. This project used ARCHER and JADE via the UK High-End Computing Consortium for Biomolecular Simulation, HECBioSim (http://hecbiosim.ac.uk).

## ABBREVIATIONS

MR: Muscarinic receptor
GPCR: G-protein coupled receptor
hMR3: human Muscarinic receptor 3
hMR2: human Muscarinic receptor 2
COPD: chronic obstructive pulmonary disease
ECL2: extracellular loop 2
TM5: transmembrane region 5
TM6: transmembrane region 6
TM7: transmembrane region 7
ICL3: intracellular loop 3
US: unbound state
BS: bound state
TS: transition state
ML: machine learning
MLTSA: machine learning transition state analysis
FEP: free energy profile
MLP: multi-layer perceptron
GBDT: gradient boosting decision tree
RAD: relative accuracy drop
RFI: relative feature importance
PCA: principal component analysis
CV: collective variable
MD: molecular dynamics;

## Notes

https://github.com/pedrojuanbj/hMR3_Unbinding

### Competing Interest Statement

The authors have declared no competing interest.

### Summary of Updates

Displaying incorrect name of first author: Pedro Juan Buigues Jorro, which should be Pedro J. Buigues

