## Supplementary Information with Tables/Figures for "Investigating the Unbinding of Muscarinic Antagonists from the Muscarinic 3 Receptor"

#### MD Simulations

##### Unbinding Trajectories

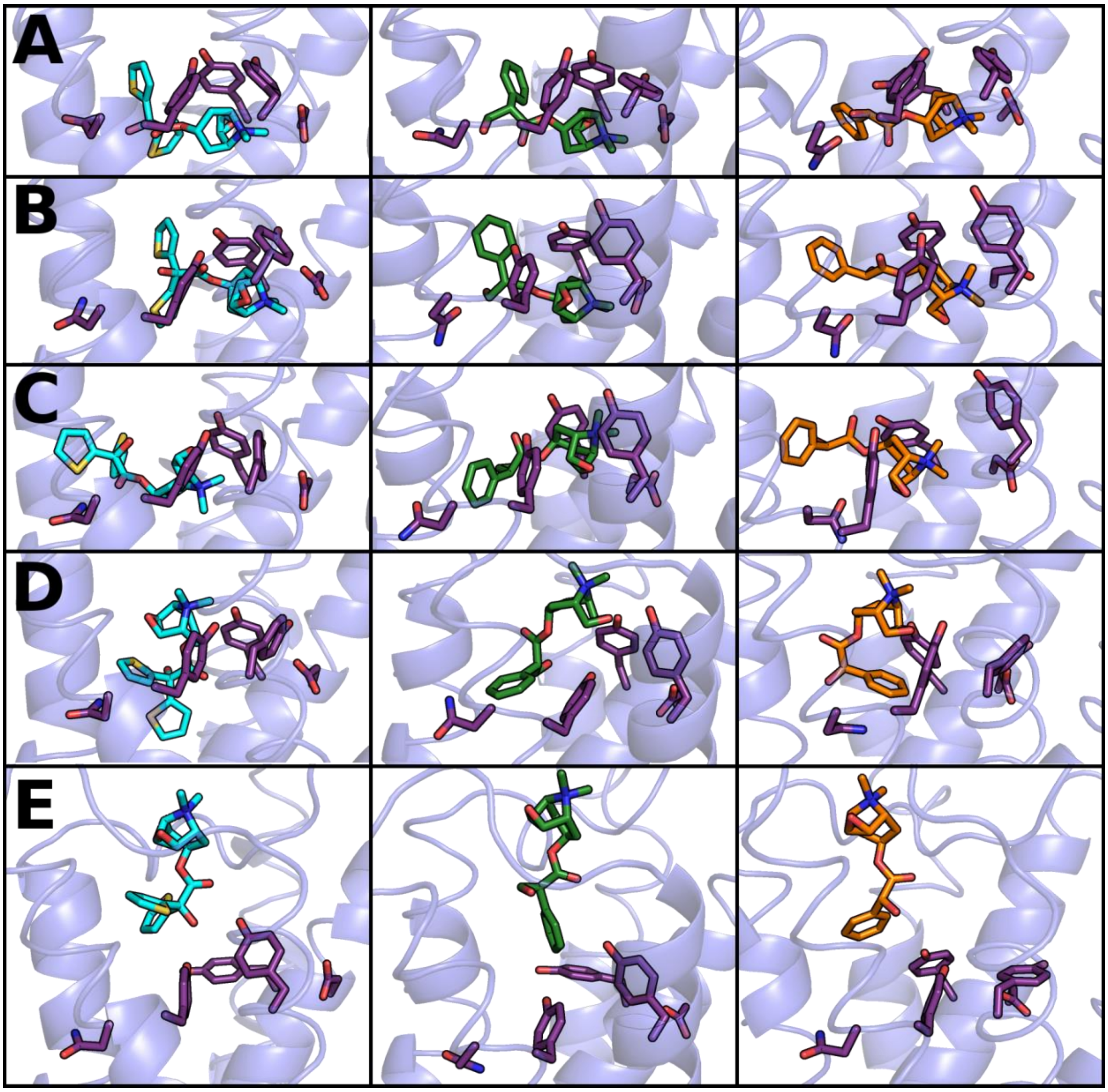

*Figure S1 – Key frames of the unbinding path of ligands 1 (left, cyan), 2 (middle, green) and 3 (right, orange) in sticks. hMR3 is represented as cartoon, residues forming the aromatic cage of the orthosteric binding site are also represented as sticks (in purple). Frames start from being bound in A, to different points in the unbinding process for each ligand, finishing with the ligand on its way to the extracellular vestibule in frame E, this is considered to be the TS of the whole process.*

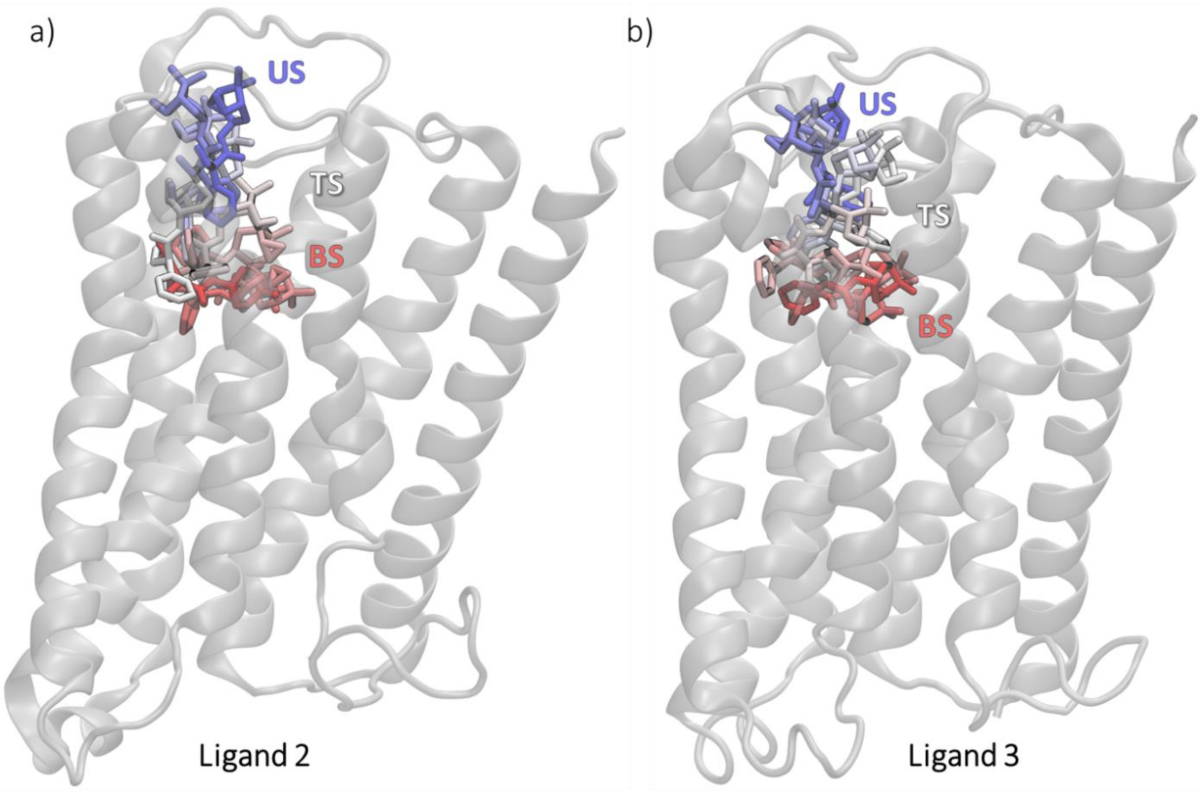

*Figure S2 – Overlay of structures found throughout the unbinding path of ligand 2 and 3 starting from hMR3's orthosteric binding site. Ligands, represented as sticks, start at the BS (red) and move through the TS (white) to reach the US (blue) on the extracellular vestibule of the receptor (grey, in cartoon).*

#### Downhill Trajectories

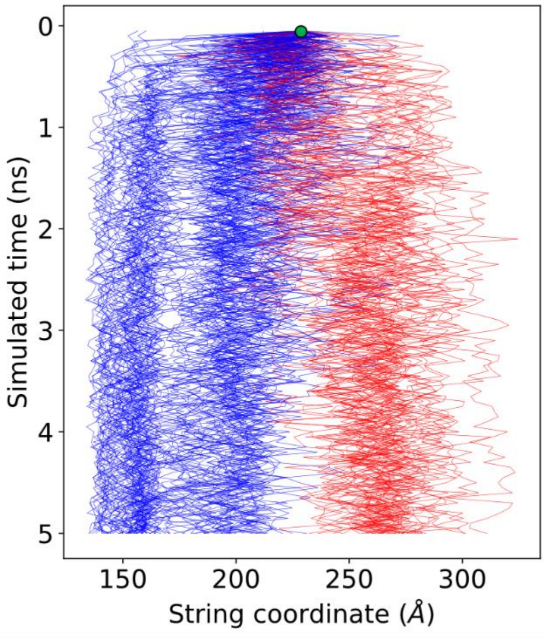

*Figure S3. Evolution of the collective variables through the downhill simulations and their classification as IN (blue) or OUT (red) for further analysis.*

### MLTSA

#### Dataset Creation

All datasets containing interatomic distances were created using mdtraj, the outcomes of the simulations were assigned using the original biasing CV value from unbinding. If after 5ns the distance was under XÅ it was assigned IN, otherwise if bigger than YÅ it was assigned OUT.

##### XYZ-PCA Dataset

To assess the importance of protein-protein distances, an additional dataset was created using the XYZ coordinates of all protein atoms (except hydrogens). The number of protein atoms for the system is ~2247, which yields around 6741 coordinates. To be able to reduce this number, a PCA projection was applied to the data. In Figure X, the resulting explained variance ratio can be found for the different PCA components resulted from the calculation.

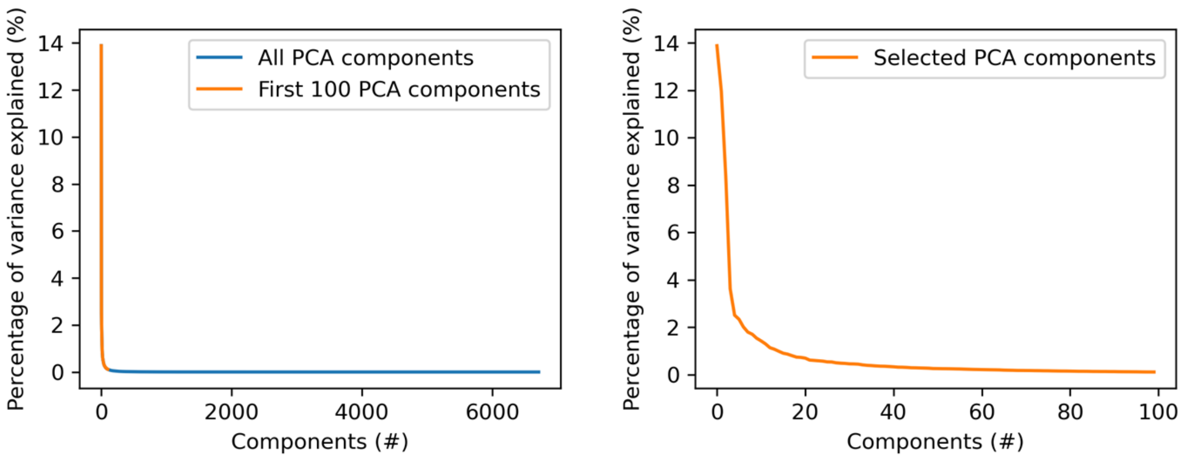

*Figure S4 – Explained variance ratio for the fitted PCA from the XYZ-PCA set. Left: Values of all PCA components projected. Right: Values more in detail for the selected components which are the first 100 components. The first 100 were selected because of the rapid decay to 0% explained variance and make sure no information was lost.*

At around 100 components, the explained variance ratio dropped close to 0. We decided to use these 100 new components as features for training.

##### 3Å and 6Å Datasets

The dataset was created by selecting all protein atoms within 3Å of any of the ligand atoms in the starting TS structure and defining all interatomic distances between ligand atoms and protein heavy atoms.

##### 3Å+Loop Datasets

To assess the importance of particular residues outside the pocket, datasets with all interatomic distances from ranging from I222 to T231 were added to the previous 3Å set, including the ECL2-TM5 junction region. (ECL2-TM5 dataset).

To validate the relevance of the ECL2-TM5 loop, an additional dataset with residues I501 to C516 around ECL3 were also tested. However, no distances from this loop were identified as important either by MLP or GBDT models.

### ML Models

#### Classifiers

##### *MLP*

The MLP model was setup using the MLPClassifier from Scikit-Learn [1], using 3 layers (input, hidden, output) with as many input nodes as features used in the first layer, 100 hidden nodes in the second layer using the ReLU[2] activation function and 1 output layer made of 1 node with a logistic activation function (0-IN,1-OUT). The model was optimized using the Adam solver,[3] with a learning rate of 0.001, iterating over data until convergence or upon reaching the maximum number of iterations (500 epochs). Convergence is determined by the tolerance and the number of epochs with no change in loss. When having 10 consecutive epochs with less than 0.0001 improvement on the loss, the training stops, and it is considered that the model has reached convergence. All datasets used the same parameters for training.

##### *GBDT*

The GBDT model used was setup using the GradientBoostingClassifier implementation from Scikit-Learn,[1] we trained 500 decision stumps as weak learners minimizing a logistic loss function, with a learning rate of 0.1. The Friedman Mean Squared Error (MSE) was used for to assess the split of the internal nodes, using a minimum of 2 samples to split, and 1 sample required at the leaf nodes. The maximum depth of the individual estimators was 3, without a limit on the maximum number of features to consider for the best split or the leaf nodes. Training was done using a validation fraction of 0.1 internally.

#### Trainings

##### *Training at different times*

To find the optimum time-range from the downhill trajectories to train the ML models, a training at different times was performed. Using a time window of 0.05 ns at a time, we tested from 0.05 ns to 0.5 ns. It was found that between 0.05 ns and 0.1 ns would be as early as possible without sacrificing reasonable accuracy.

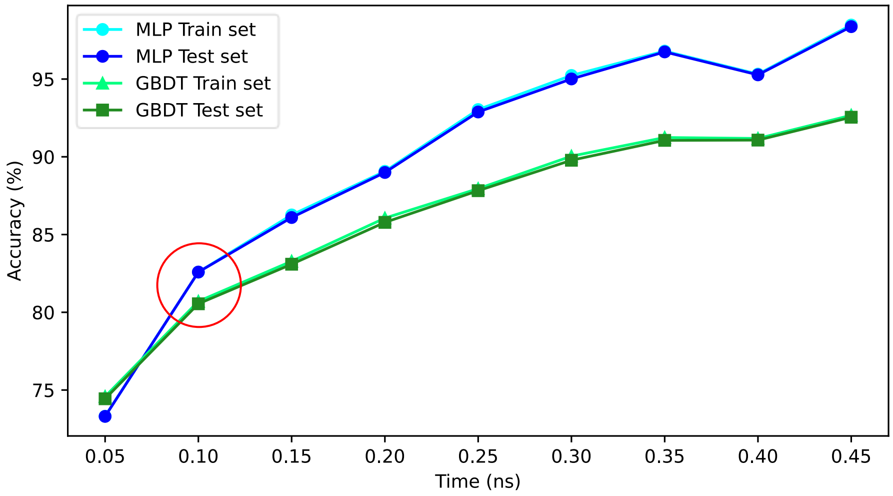

*Figure S5 – MLP (Blue) and GBDT (Green) accuracy for the test and train sets at different times throughout the downhill trajectories using early near-TS data until 0.5 ns. Highlighted in red, the chosen time-frame to use in further trainings, from 0.05 to 0.10 ns. Note the dataset used is the 3Å set.*

*Table S1 – Train (70% data) and test (30% data) accuracy for the MLP and GBDT models on the different datasets.*

| | | MLP (%) | GBDT (%) |
|---|---|---|---|
| 3Å set | *Train* | 77.80 ± 0.32 | 76.15 ± 0.75 |
| | *Test* | 76.34 ± 0.52 | 75.89 ± 0.74 |
| 6Å set | *Train* | 75.18 ± 0.23 | 76.64 ± 0.12 |
| | *Test* | 75.71 ± 0.25 | 76.31 ± 0.11 |
| 3Å+ECL2 set | *Train* | 76.82 ± 0.95 | 80.31 ± 0.15 |
| | *Test* | 76.27 ± 1.14 | 80.14 ± 0.12 |
| XYZ-PCA set | *Train* | 100 | 93.42 ± 0.82 |
| | *Test* | 100 | 93.25 ± 0.85 |
| Allres set | *Train* | 77.85 ± 0.01 | 79.27 ± 0.01 |
| | *Test* | 77.71 ± 0.01 | 79.01 ± 0.01 |
| Allres+wat set | *Train* | 81.27 ± 0.04 | 80.22 ± 0.03 |
| | *Test* | 81.05 ± 0.04 | 79.82 ± 0.03 |

#### Feature Selection

##### XYZ-PCA Analysis

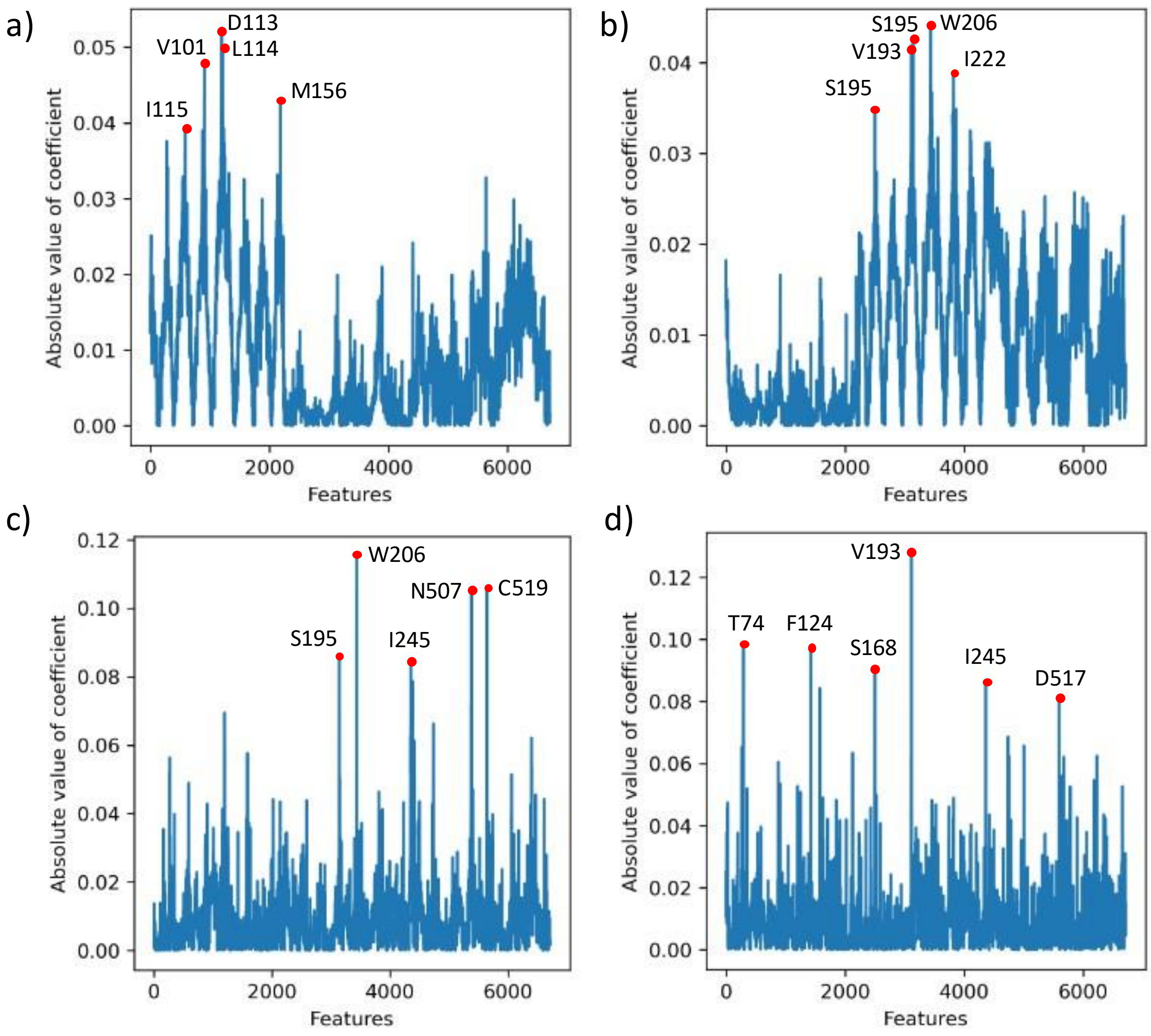

*Figure S6. Atomic xyz feature contributions to the PCA components 1 (b), 2 (b), 23 (c), and 59 (d).*

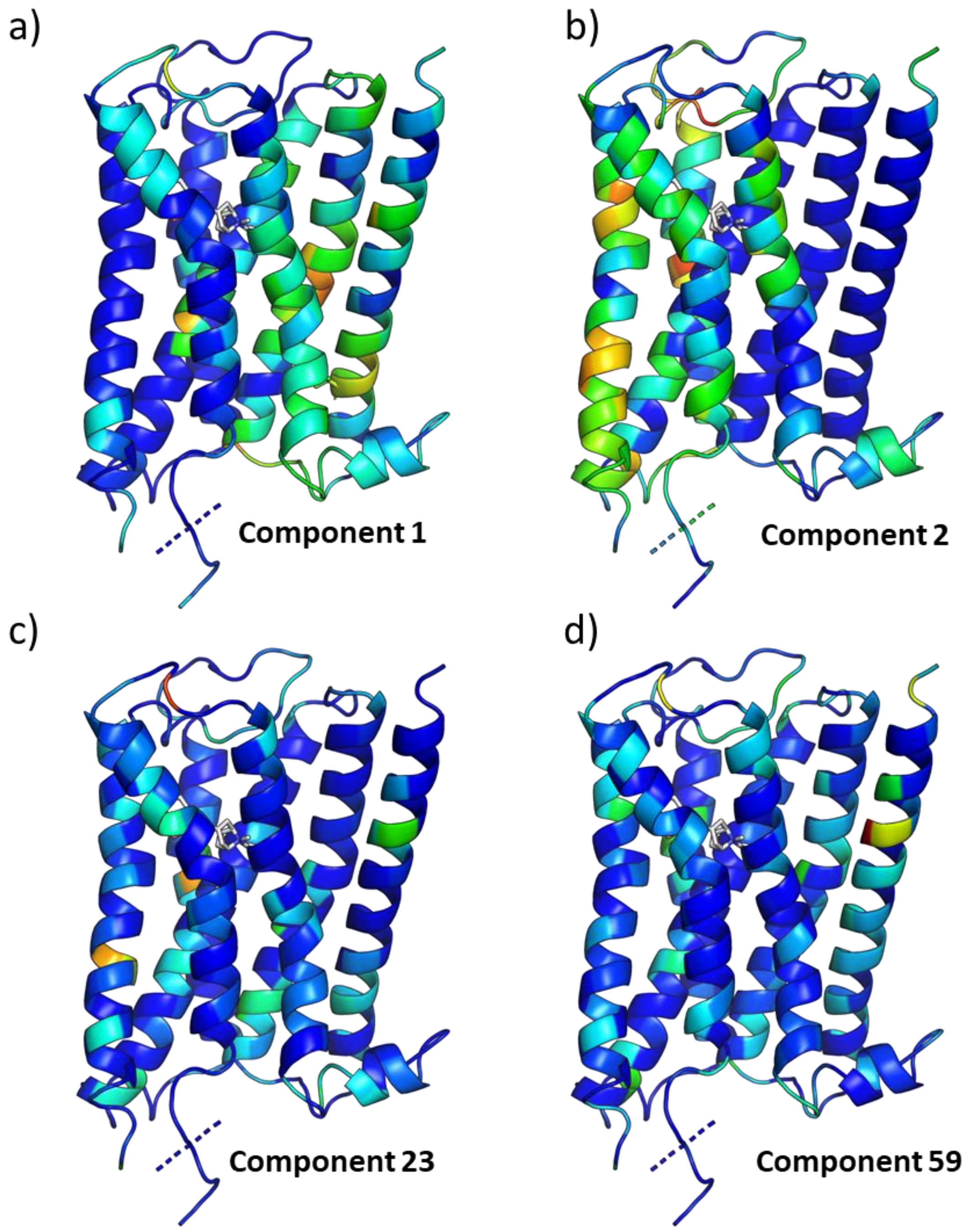

*Figure S7. Average atomic contributions to the PCA components 1 (b), 2 (b), 23 (c), and 59 (d) colored by the R-factor (blue to red: zero to 1.*

#### 6A Dataset

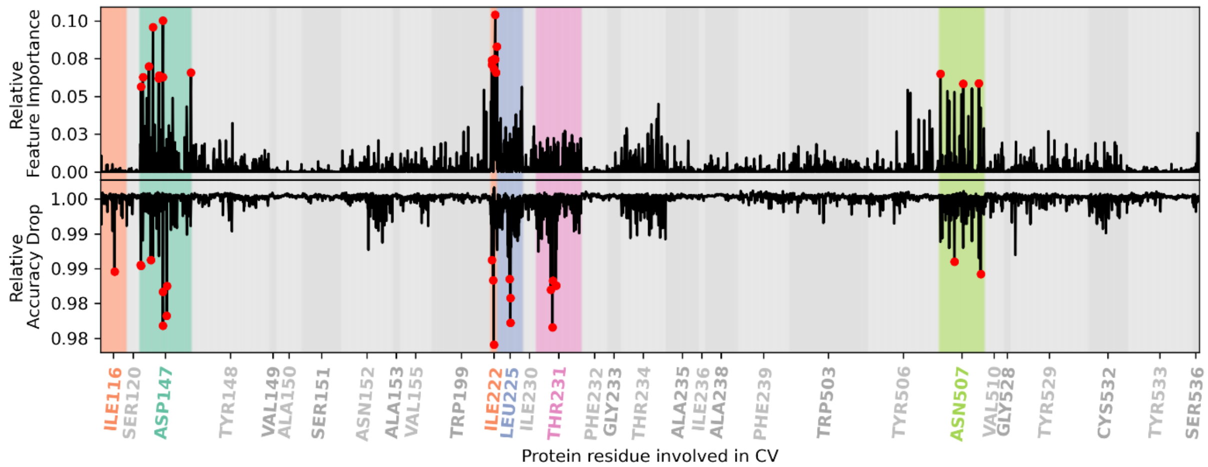

*Figure S8 – RAD and RFI for individual interatomic distances for the 6 Å dataset. Top distances marked in red, and top residues highlighted in color.*

#### 3A+ECL3 Loop

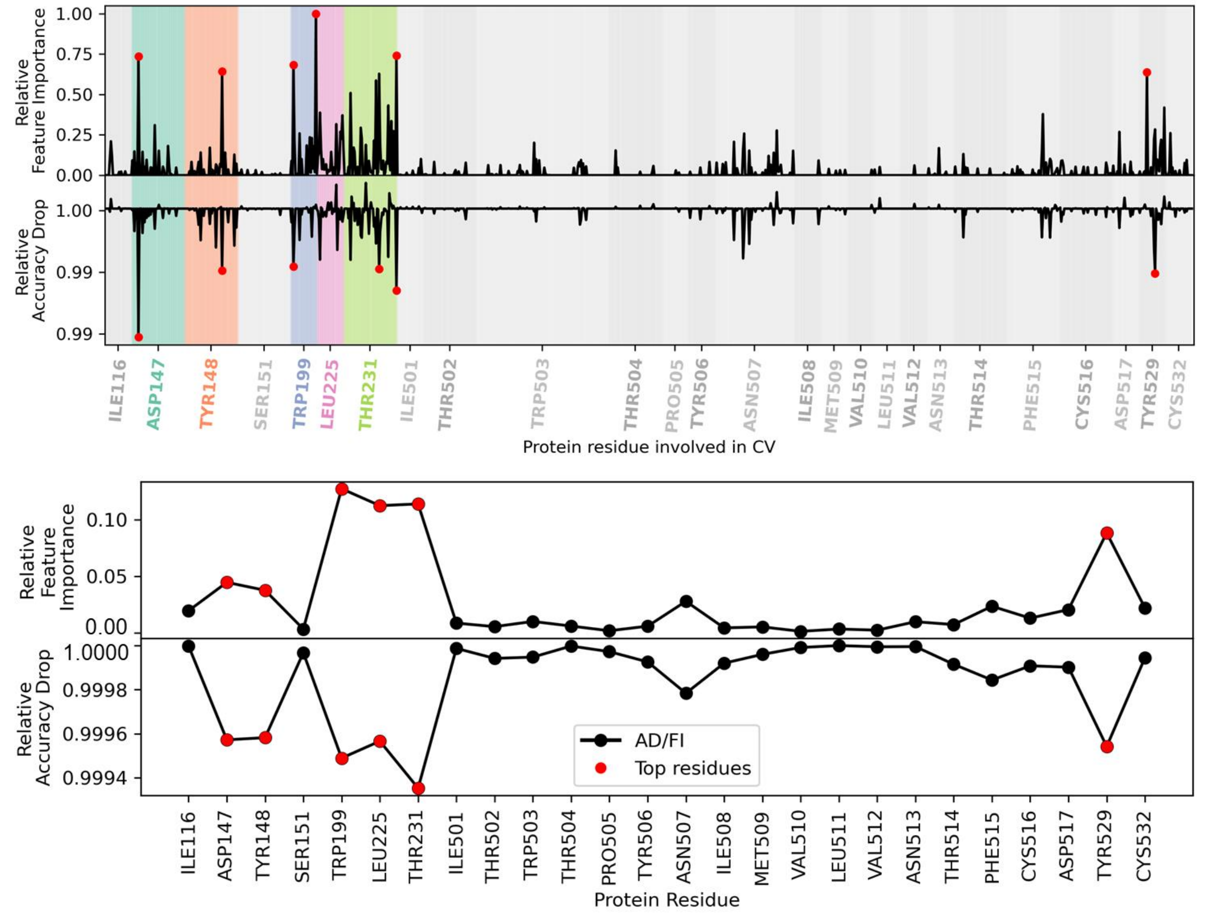

*Figure S9 – Top: RAD and RFI for the 3Å* adding interatomic distances from an alternate loop ECL3. Residues newly included range from I501 to C516. Top distances marked in red, top residues highlighted in color. Bottom: Average per residue RAD and RFI for the 3Å+ECL3. Top residues marked in red.

#### Additional resources

A github repository with the multi-PDB and GIF unbinding trajectories through the string windows and the PDB TS structures can be found at: https://github.com/pedrojuanbj/hMR3_Unbinding
